# Multivariate State Hidden Markov Models for Mark-Recapture Data

**DOI:** 10.1101/025569

**Authors:** Devin S. Johnson, Jeff L. Laake, Sharon R. Melin, Robert L. DeLong

**Affiliations:** National Marine Mammal Laboratory, Alaska Fisheries Science Center, NOAA

**Keywords:** Capture-recapture, Cormack-Jolly-Seber, Hidden Markov Model, Multivariate, Partial observation, State uncertainty

## Abstract

State-based Cormack-Jolly-Seber (CJS) models have become an often used method for assessing states or conditions of free-ranging animals through time. Although originally envisioned to account for differences in survival and observation processes when animals are moving though various geographical strata, the model has evolved to model vital rates in different life-history or diseased states. We further extend this useful class of models to the case of multivariate state data. Researchers can record values of several different states of interest; e.g., geographic location and reproductive state. Traditionally, these would be aggregated into one state with a single probability of state uncertainty. However, by modeling states as a multivariate vector, one can account for partial knowledge of the vector as well as dependence between the state variables in a parsimonious way. A hidden Markov model (HMM) formulation allows straightforward maximum likelihood inference. The proposed HMM models are demonstrated with a case study using data from a California sea lion vital rates study.

## 1. INTRODUCTION

The seminal papers by Cormack (1964), Jolly (1965), and Seber (1965) initiated 50 years of active development of capture-recapture theory and application. The Cormack-Jolly-Seber (CJS) model for survival estimation and the Jolly-Seber model for survival and abundance estimation and their extensions are still the most widely used capture-recapture models. They have been implemented in the computer software `MARK` (White and Burnham, 1999), `POPAN` (Arnason and Schwarz, 1999), `SURGE` (Lebreton et al., 1992), `M-SURGE` (Choquet et al., 2004), `E-SURGE` (Choquet, Rouan and Pradel, 2009), and, more recently, `marked` (Laake, Johnson and Conn, 2013) and `multimark` (McClintock, 2015).

Expansion of CJS models to account for movement between areas (multistate) was initiated by Darroch (1958) and Arnason (1973) and further developed by Hilborn (1990), Hestbeck, Nichols and Malecki (1991), Schwarz, Schweigert and Arnason (1993) and Brownie et al. (1993). Nichols et al. (1992) used the multistate model to describe survival with state transitions based on mass and this has been followed by numerous other examples using the multistate model (Lebreton and Pradel, 2002; Nichols and Kendall, 1995; White, Kendall and Barker, 2006). An important advance to the multistate CJS generalization was made by Kendall et al. (2004) who considered models with state uncertainty (unknown state) and Pradel (2005) who cast a multistate model with state uncertainty as a hidden Markov model (Zucchini and MacDonald, 2009). The state uncertainty models are often termed multievent models. In a further step, Kendall et al. (2012) combined a robust design with a multistate hidden Markov model to improve precision in the face of state uncertainty. We use the general term *state-based* to refer to all of these types of CJS models.

For traditional CJS analysis, the detection portion of the model is often considered to be a nuisance with little scientific interest and often determined largely by the resampling methods used by researchers to resight individuals. The achievement made by the development of the original CJS model was that it permitted estimation of survival even when individuals were not observed on every occasion. The state-based extensions allowed survival inference to account for time-varying heterogeneity or deviations from CJS model assumptions induced by states as well make inference to the state process itself (Lebreton and Pradel, 2002). With the recently developed extensions, one can often find state-based CJS analyses where survival is also considered a nuisance process and primary scientific interest lies with the transitions between states. See Gourlay-Larour et al. (2014) for a recent example. With this in mind, practicing ecologists are using the state-based CJS framework and the associated HMM formulation to analyze complex state transitions such as the combination of geographic location and reproductive or disease status. This presents the problem that a state may be only partially observed when an individual is resighted. For example, location will be known but reproductive status may not be observed. To further generalize the state-based CJS models, King and McCrea (2014) developed a closed form likelihood expression to handle partial knowledge of the state of an individual. However, a general framework for parametrizing complex state-based CJS models is not examined. Laake et al. (2014) made initial strides into multivariate state CJS models by proposing a bivariate state model to account for dependence in double-mark loss. Here we seek to augment the work of King and McCrea (2014) and Laake et al. (2014) to provide a general method to construct complex state-based CJS models.

In this paper, we propose a general modeling framework for multivariate state-based CJS models in which the state is defined by one or more discrete categorical variables and each variable may be unknown when the animal is resighted. The modeling framework extends the state-based models in MARK by allowing any or all of the state variables to be uncertain rather than just a single uncertain state. We have implemented the model in the `marked` package (Laake, Johnson and Conn, 2013) for the R statistical environment (R Development Core Team, 2015) using a hidden Markov model (HMM) formulation for maximum likelihood estimation of the parameters. General multivariate HMMs have been proposed in the past (Ghahramani and Jordan, 1997; Brand, Oliver and Pentland, 1997), but here we model the state vector in a log-linear framework (Christensen, 1997) to easily allow all levels of dependence among state variables. The proposed model is illustrated with a case study using 18 years of annual resighting data collected on a single cohort of California sea lion *(Zalophus californianus)* pups that were both branded and double tagged on their flippers in 1996 at San Miguel Island near Santa Barbara, California (Melin et al., 2011). The animal’s state was defined based on location (San Miguel or Año Nuevo Islands) which was always known when the animal was resighted and the, possibly unknown, states of the two flipper tags (present or absent due to tag loss).

## 2. METHODS

To begin the description of multivariate state modeling of mark-recapture data, we must begin with the state and data structure. The general structure is based on the capture histories of individuals, indexed *i* = 1, …, *I*, over capture occasions, indexed *j* = 1, …, *J*. The state of an individual on a particular occasion is determined by the cross-classification of that individual to a cell in a multiway table where the margins represent state variables of interest. Specifically, the state, s = (*s*_1_, …, *s_K_*) is a vector where the entries correspond to *K* classification criteria, e.g., location, reproductive status, or disease presence. Each state variable has a set of possible values 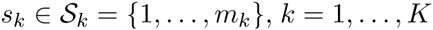. The space of the state vector, s, is thus 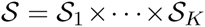. The number of possible states is denoted 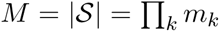. We also need to augment the state-space, 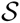, with a “death” state, such that the individual can transition between all 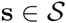 as well as death. We represent the death state with s = 0, so the augmented state space is 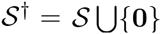. The observed data for an individual on a particular (re)capture occasion is denoted as c = (*c*_1_, …, *c_k_*). The observed state can differ from the true state in two ways. First, an individual may not be observed (detected) on a particular capture occasion. Second, it is possible, and highly probable in many situations, that even if an individual is physically observed, state specification may not be completely determined by the researcher. Thus, the space of the observation vector is augmented with two additional levels corresponding to “not observed” and “unknown,”, i.e., *c_k_* ∈ {0,1, …, *m_k_*, *m_k_* + 1}, where 0 is for undetected individuals and *m_k_* + 1 is associated with an unknown level for that state variable. In heuristic descriptions we will use *c_k_* = *u* to be clear we are referring to the unknown state. In addition, we take the convention that if = 0 for any k, it is assumed that all *c_k_* = 0. We use *c_ij_* to denote the observed state of individual *i* on occasion *j*. Table A.1 of Supplement A (Johnson et al., 2016a) provides a glossary of the notation used in this paper.

### 2.1 Two state variable model

Specifying a model for a general multivariate discrete state is notationally cumbersome due to the complex dependence structure. Therefore, we begin with a description for a bivariate state model, and then follow up in the next section with a general model formulation for > 2 state variables with additional covariates. The general parameter types are the same as in any state-based CJS model: survival (*ϕ*), detection (*p*), and state transition (*ψ*) from one occasion to the next. In addition, we have parameters associated with the inability to fully assess the state of an individual (*δ*) and the probability that an individual is in a particular state at first capture (*π*). Typically, each of these parameter types depends on the state of the individual; e.g., survival of individual *i* from occasion *j* to occasion *j* + 1 depends on the state of the individual on occasion *j*. Each multivariate state variable may contribute independently to survival, or may interact at various levels to further enhance or degrade survival. Thus, models for these parameters must accommodate this interaction (dependence) when modeling state transition. We take a log-linear approach (Christensen, 1997) for modeling multiway contingency tables which allows for all levels of dependence between state variables.

We begin with modeling state transitions from one occasion to the next. Suppose individual *i* is alive on occasion *j* < *J*, then the probability of a transition from state 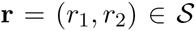 on occasion *j* to state 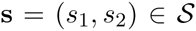 on occasion *j* + 1 is given by the conditional log-linear model,

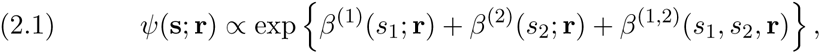

where, for each 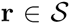, *β*^(1)^(*s*_1_; r), *β*^(2)^(*s*_2_; r), and *β*^(1,2)^(*s*_1_, *s*_2_; r) are parameters which depend on the previous state, r, and are indexed by the values of *s*_1_, *s*_2_, and the double index of (*s*_1_, *s*_2_), respectively. For example, the collection of *β*^(1)^ parameters is {*β*^(1)^(1; r), …, *β*^(1)^(*m*_1_; r)}. If we denote to be the transition matrix describing the probability of transitioning from any state on occasion *j* to any other state on occasion *j* + 1, equation (2.1) evaluated for all 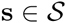would form the rows associated with each possible current state r. There are *m*_1_ + *m*_2_ + *m*_1_*m*_2_ parameters for each r. Thus, there are *m*_1_*m*_2_ × (*m*_1_ + *m*_2_ + *m*_1_*m*_2_) *β* parameters within the *m*_1_*m*_2_ × *m*_1_*m*_2_ transition matrix Ψ. Hence, the model is overparameterized. To alleviate this problem it is customary to pick a reference cell say, 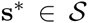, and fix all *β* parameters to 0 when 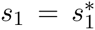 *o*r 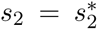. For example, if s* = (*r*_1_,*r*_2_) (the current state), *β*^(1)^(*r*_1_;r) ≡ 0, *β*^(2)^(*r*_2_; r) ≡ 0, *β*^(1,2)^(*s*_1_,*r*_2_; r) ≡ 0 for any *s*_1_ value, and *β*^(1,2)^(*r*_1_,*s*_2_; r) ≡ 0 for any *s*_2_ value. Of course it is not necessary to use the current state as the reference state, but it has some advantages, namely, the *β* parameters are interpreted as controlling movement away from the current state and the current state is usually an acceptable possibility for a realization at the next occasion (i.e., non-zero probability of occurring). This may not be the case if you define a fixed state as the reference (see Section 3 for example). See Table (1) for an example of *ψ* (s; r) specification with the reference cell constraint.

**Table 1.** *Example transition probabilities, ψ*(s; r), *for a hypothetical bivariate state* s *composed of state variables s*_1_ ∈ {1, 2, 3} *and s*_2_ ∈ {1, 2}. *Here we use the reference cell* s* = (1,1). *The p probabilities are presented in their unnormalized form as in equation (2.1). The necessary normalizing constant is the sum over all the table entries.*

| | | $s_2$ | |
| --- | --- | --- | --- |
|  |  | 1 | 2 |
| $s_1$ | 1 | 1 | $\exp\{\beta^{(2)}(2; \mathbf{r})\}$ |
| | 2 | $\exp\{\beta^{(1)}(2; \mathbf{r})\}$ | $\exp\{\beta^{(1)}(2; \mathbf{r}) + \beta^{(2)}(2; \mathbf{r}) + \beta^{(1,2)}(2, 2; \mathbf{r})\}$ |
| | 3 | $\exp\{\beta^{(1)}(3; \mathbf{r})\}$ | $\exp\{\beta^{(1)}(3; \mathbf{r}) + \beta^{(2)}(2; \mathbf{r}) + \beta^{(1,2)}(3, 2; \mathbf{r})\}$ |

Markovian dependence in time is explicitly included by allowing the *β* parameters to depend on the previous state. In addition, dependence between state variables is accommodated via the *β*^(1,2)^ interaction parameters. If *β*^(1,2)^ (*s*_1_, *s*_2_; r) = 0 for all values of *s*_1_ and *s*_2_ then this is a necessary and sufficient condition for *s*_1_ and *s*_2_ to be independent given the previous state r. Sufficiency is easily observed by the fact that under that assumption

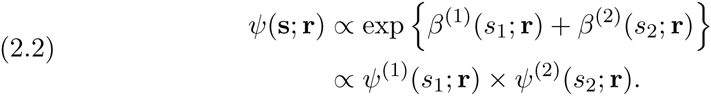

Thus, the probability of transition to state s is the product of independent transitions from *r*_1_ → *s*_1_ and *r*_2_ → *s*_2_. Proof of necessity is quite cumbersome and beyond the scope of this paper. See Lauritzen (1996) for more theoretical details of log-linear modeling. One caveat of these independence results is that they assume there are no structural zeros (impossible state transitions). If there are, one should check that independence still holds if it is a desirable property. Otherwise, zero valued interaction terms can just be used for parsimonious modeling.

The next set of parameters of scientific interest is survival. Given that individual *i* is in state s on occasion *j* the probability of survival from occasion *j* to *j* + 1 is described by

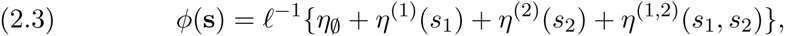

where *ℓ* is a link function such as logit or probit, and the *η* parameters are functions of (indexed by) the current state, s. The *η* parameters have the same structure and reference cell constraints as the *β* parameters in the *ψ* model. Readers should also observe that there is a 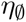 parameter. This term is not indexed by the state and represents baseline survival. Even though the notation may appear different for some readers, it should also be noted that this *ϕ* model is identical to a model with *s*_1_ and *s*_2_ being thought of as known categorical covariates with interaction effects included. So, as far as the survival model is concerned, *s*_1_ and *s*_2_ are equivalent to (individual × occasion) indexed factor covariates.

We now move to the observation and detection portion of the model. Given that individual *i* is in state s on occasion *j*, the probability that the individual is detected (i.e., recaptured or resighted) by the researcher is given by

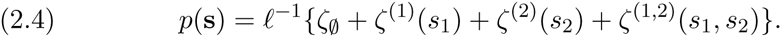

As with the survival model, the state variables can be considered as factor variables for the purposes of conceptualizing the model. There is one more set of parameters working in concert with *p*, these are related to the ability of the researcher to jointly classify a detected individual according to all of the state variables. If an individual *i* is detected on occasion *j*, then the joint probability of the state observation, c = (*c*_1_, *c*_2_)^′^, conditioned on the true state is modeled via,

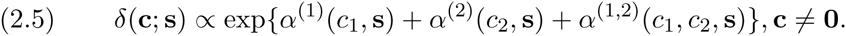

Note that we do not consider misidentification of states; a state variable is either observed correctly (*c_k_* = *s_k_*) or it is not observed and the value is unknown (*c_k_* = *m_k_* + 1 or *u*). As with the multivariate transition model, *ψ*, a reference cell, say c* is necessary for parameter identifiability. One could use c* = (*m*_1_ + 1, *m*_2_ + 1), the completely unknown state, to be consistent with previous univariate state uncertainty models (e.g., Laake 2013; Kendall et al. 2012), but it is not necessary and may be undesirable if the completely unknown state is not observable; e.g., double-tag studies where no third permanent mark is available. We suggest using the current state, c* = s, as the reference. As with the *ψ* models, one can incorporate dependence in recording the level of each state variable. That dependence implies, for example, that if the researcher is able to record the level for one state variable, the researcher is more (less) likely to record the level of the other. There may be some state variables which are always recorded with certainty, e.g., location. So, for any c that contains *c_k_* = *m_k_* +1 for those state variables, we can also fix the *δ* ≡ 0 and remove the corresponding *α* parameters from the overall parameter set. As with the state transition probabilities, *ψ*, if these structural zeros exist, then the dependence interpretation needs to be investigated because conditional dependence can be induced by the zeros.

Finally, the last set of parameters is associated with the initial capture of the individual. We are considering only CJS type models here, thus, we are still conditioning the model on the occasion when an individual is first observed. When an individual is first captured (marked), however, the researcher may not be able to observe all state variables. Therefore, we need to model the probability that individual *i*, who is first marked on occasion *j*, is in state s. So, we describe this probability via,

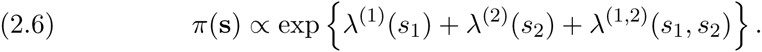

Here, there is no previous cell from which to form the reference cell. Therefore the researcher will have to decide on the most appropriate state to serve as the reference. The *π* model parameters, however, may not necessarily have to be estimated. For example, if it is certain that all state variables will be observed at first marking then it can be fixed to *π*(s) = 1, where s is the state individual *i* was in when it was first marked, and *π*(r) = 0 for all r ≠ s (see Section 3). Or, a researcher might specify, say, *π*(s) = 1/*M*, where *M* is the number of cells in 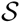. The *π* probabilities can be interpreted as a prior distribution, in the Bayesian sense, on the state of a newly marked individual, thus, 1/*M* would be a uniform prior over the state-space.

### 2.2 General multivariate state model

In this section we generalize the bivariate state models presented in the previous section. Here, researchers can consider state specifications that index a cell in a general *K* dimensional hypercube and models can depend on covariates, as well. To accomplish this we introduce the notation 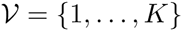 to be the set which indexes the collection of *K* state variables and 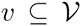 is a subset. The partial state s^(^*^v^*^)^ represents only those elements of s whose indexes are contained in 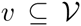. The state subspace 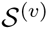 denotes the set of all possible values of s^(^*^v^*^)^.

Now, the full specified model describing the transition of individual *i* in state r = (*r*_1_, …, *r_K_*) on occasion *j* to state s = (*s*_1_, …, *s_K_*) on occasion *j* + 1 is given by,

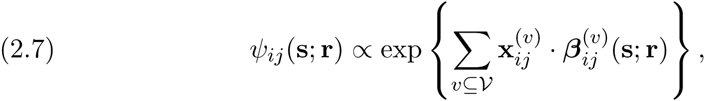

where 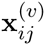 is a vector of covariates, the notation 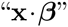 represents the dot product 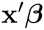, and 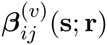 is a coefficient vector that depends on the state, 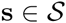, only through the value of the sub-state 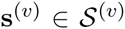 and 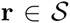. As with the bivariate model in the previous section, we must satisfy the reference cell constraint to have an identifiable set of parameters. Therefore, all *β*^(^*^v^*^)^(s; r) ≡ 0 if any element of the sub-state s^(^*^v^*^)^ is equal to the corresponding elements of the reference cell, s*. Independence of state variables is more complex than described in the previous section due to a number of possible interactions between different sets of variables. To provide a generalization, let *v*_1_, *v*_2_, and *v*_3_ be subsets of state variables that partition 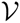. Then 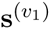 and 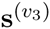 are conditionally independent given 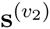 (and r) if and only if *β*^(^*^v^*^)^(s; r) = 0 for all s, where *v* contains elements of both *v*_1_ and *v*_3_ (i.e., 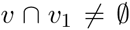 and 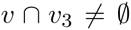) (Frydenberg, 1990). For example, suppose 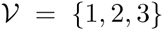, then *s*_1_ is conditionally independent of *s*_3_ given *s*_2_ (and r) if *β*^(1,3)^(s; r) = *β*^(1,2,3)^ (s; r) = 0 for all s. Further, if we assume that *β*^(1,2)^(s; r) = 0, then s^(1)^ = *s*_1_ is independent (given r) of s^(2,3)^ = (*s*_2_,*s*_3_). In that case, we can follow the example in (2.2) and write

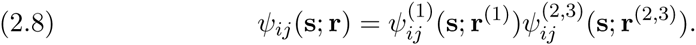

This independence factorization can extend to more than two groups using the Gibbs factorization theorem (see Frydenberg 1990). Again, independence theorems are based on the absence of structural zeros.

We can progress through the other parameter groups in a similar fashion to produce a general multivariate state model. The survival portion of the model can be generalized to

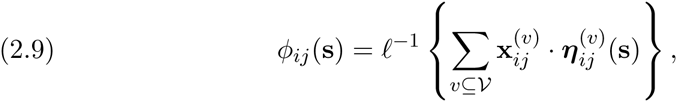

where the covariate vector 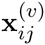 need not be the same as that shown in equation (2.7); we use the same notation simply to avoid extra clutter. Cycling through the remaining parameters, we have:

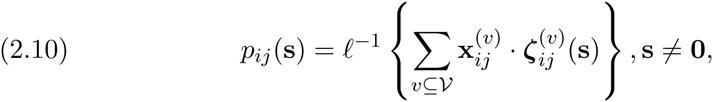

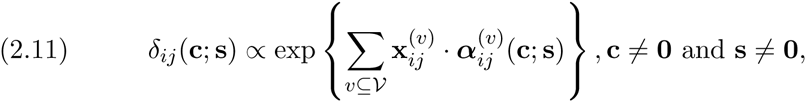

where we suggest the reference cell c* = s. In general, there will be 2*^d^* possible values of c associated with each s, where *d* is the number of state variables for which a *u* can be recorded. Finally, the probability distribution of states upon first capture is

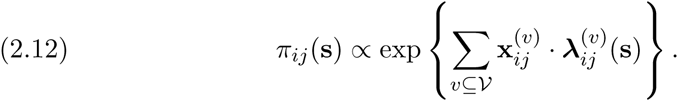

This completes the general model for a *K* dimensional state vector. In the next section we focus on methods for efficient parameter estimation and inference. Similar to the *ψ* models, the *δ* and *π* parameters can be factored into independent products as well. This will facilitate easier overall model construction if certain state variables can be assumed to be independent, making it easier to parameterize dependence over a small subset of variables.

### 2.3 Statistical inference

Statistical inference for multistage mark-recapture models can be challenging (McCrea and Morgan, 2014). We take the approach described by Laake (2013) who uses a hidden Markov model (HMM) formulation. By framing a state-based CJS model as a specific HMM the efficient *forward algorithm* can be used to maximize the likelihood (Zucchini and MacDonald, 2009). Laake (2013) provides a detailed description of HMM formulated mark-recapture models and likelihood calculation, but we briefly revisit it here to place the forward algorithm in the context of multivariate state CJS models.

To begin the description of the forward algorithm in the multivariate state CJS model case, we must define three matrices. The transition probability matrix for movement within 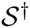 is given by,

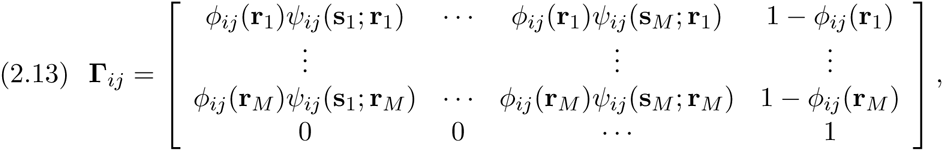

Where *M* is the size of the state-space (recall *M* = Π*_k_ m_k_*), r*_m_*, *m* = 1, …, *M*, refers to the current state on occasion *j*, s*_m_* refers to the state to which the animal transitioning on occasion *j* + 1, and the last column (row) is associated with transition to (from) the death state. In addition to the state-space augmentation, we define the observation probability matrix which describes the probability of observing state c*_n_*, *n* = 0, …, *N*, for animal *i* on occasion *j* (rows) given it was actually in state 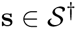 (columns),

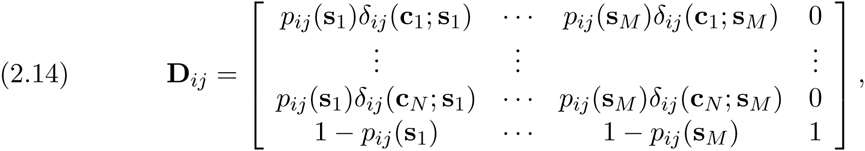

where observed state c_0_= (0, …, 0), the undetected state, is represented by the last row. For use in the forward algorithm, we also define the *K* × *K* matrix, P(c*_ij_*) to be a diagonal matrix with the row of D*_ij_* associated with observed state for individual *i* on occasion *j* as the diagonal entries.

Using the Γ*_ij_* and P(*c_ij_*) matrices we can describe the forward algorithm specific to maximum likelihood estimation in the multivariate state MR model. The algorithm proceeds as follows for individual *i* = 1, …, *I*,

1. Initial conditions:

- Set 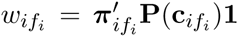 where *f_i_* = 1, …, *J* − 1, is the occasion on which individual *i* was first captured/marked, 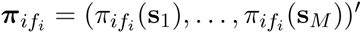, and 1 is a vector of all 1s.
- Set 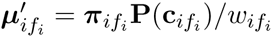
- Set 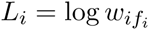
2. For *j* = *f_i_* + 1, …, *J*:

- Set 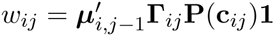
- Set 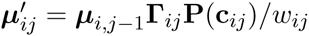
- Set *L_i_* = *L_i_* + log *w_ij_*.

The forward probabilities, *μ_ij_*· represent the conditional probability distribution of the state of individual *i* on occasion *j*, given the observed states up to and including occasion *j* and the parameters. After the algorithm has been run for every individual, the log-likelihood is given by,

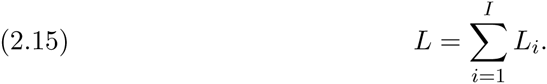

The log-likelihood can be maximized to provide parameter estimates and biological inference. In the next section we will provide a real-world example where multivariate states are collected in mark-resight study of California sea lions.

## 3. EXAMPLE: MOVEMENT, SURVIVAL, AND MARK LOSS IN CALIFORNIA SEA LIONS

### 3.1 Data and Model Description

As an example of a multivariate state CJS analysis we use 18 years of annual resighting data collected on a single cohort of California sea lion *(Zalophus californianus)* pups. In 1996, a total of 485 4-5 month old pups born on San Miguel Island (SMI), off the west coast near Santa Barbara, California (Melin et al., 2011) were herded into a pen over a total period of five days. The sex and weight of each pup was recorded and a unique permanent number was applied as hot brand to their left side. In addition, a uniquely numbered yellow roto-tag was applied to each of their foreflippers. Pups were resighted by their brand during a three month period from 15 May to 15 August each year at SMI and Año Nuevo Island (ANI). SMI is a breeding rookery, whereas ANI is a haul out where animals rest, and only a few pups are born and raised. When the animal was resighted the presence (+) or absence (−) of each tag was noted. In many cases, however, the status was not recorded for one or both of the tags. Although the flipper tags were not directly relevant to the resighting effort, their presence allowed a rare opportunity to assess loss and dependence for other pinniped studies that rely solely on flipper tags for marking (e.g., Testa et al. 2013).

From the sea lion data, a capture history was constructed with annual occasions and each entry was composed of 3 characters, with the first for the area, the second for the left tag and the third for the right tag. There were 8 different possible states for a live animal (*a* + +, *a* − +, *a* + −, *a* − −, *s* + +, *s* − +, *s* + −, and *s* − −). Here we are using symbolic representation of the states for ease of discussion. In terms of the notation presented in the previous sections, *s*_1_ is the location state (*s*_1_ = 1 for “*a*” and 2 for “*s*”), *s*_2_ and *s*_3_ are the left and right tag status variables (*s_k_* = 1 for “+” and 2 for “−”; *k* = 2, 3).

To begin the model specification used to analyze these data we will start with the observation portions of the model (*p* and *δ*). The possible capture history values were 0 if not seen, the 8 possible states if seen and each tag status was recorded, and an additional 10 observations with unknown tag status (*a* + *u*, *au* +, *a* − *u*, *au*−, *auu*, *s* + *u*, *su*+, *s* − *u*, *su*−, *suu*). For these data, location is always known with certainty when the animal is detected but each tag status could be unknown. For the 5 model we used the fully known tag status as the reference cell which implies that the parameters can be interpreted as controlling whether tag status was not obtained (i.e., unknown). The following formulation was used,

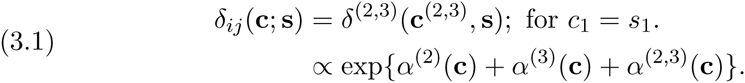

Table A.2 in Supplement A (Johnson et al., 2016a) illustrates the observation model parameters. For detection, we used the occasion × area model

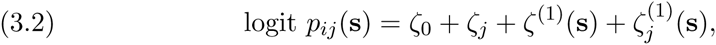

where *ζ_j_* is an occasion effect, *ζ*^(1)^(s) is an area effect, and 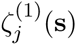 is an occasion × area interaction. Resighting of animals was based on the permanent brand mark, thus, tag status was not included in the model.

Now, we describe the more scientifically relevant, process portions of the model. First, all animals started on SMI with two tags (i.e., s*_i_*_1_ = (*s* + +)) on the first release occasion. Thus, for this example *π_i_*_1_(s) = 1 for s = (*s* + +). The survival model was formulated with sex, age, and area specific effects,

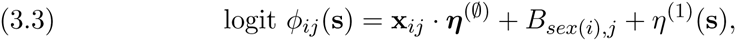

where x*_ij_* = (1, *w_ij_*), where *w_ij_* is the mass anomaly (difference from sex specific mean) if *j* = 1, zero elsewise (i.e., only pup survival is influenced by mass), *B_sex(i),j_* is a sex specific *b*-spline smooth (df = 3; Hastie 1992) over age, and *η*^(1)^(s) is an area effect. Here, mass anomaly was used as a proxy for body condition. Males tend to be heavier on average and we did not want sex differences due to other factors to be hidden within an absolute mass effect. For the state transitions we used the previous state as the reference state for the following occasion, i.e., s* = r for transitions from r → s. Thus, the parameters control movement away from the current area (*s*_1_) and loss of flipper tags (*s*_2_, *s*_3_). The state transitions were modeled by

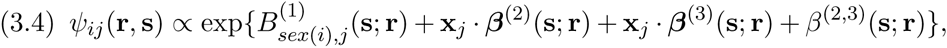

where 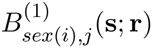 is a sex and area specific *b*-spline (df=3) over age, x*_j_* = (1, *age_j_*), and *β*^(2,3)^(s; r) is an interaction term between the two tags which models dependence in loss. Although not immediately obvious, the *b*-spline effect, 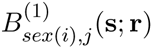, can be written in the form 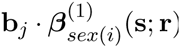, where b*_j_*· is a vector of the *b*-spline basis function values for occasion *j*. Therefore, it fits into the proposed log-linear framework. Also, because the same type of tag was placed on each flipper, we constrained the marginal loss rates to be identical for each side. This is accomplished by setting the tag loss parameters *β*^(2)^(−; +) = *β*^(3)^(−; +). The probabilities for impossible transitions (i.e., going from “−” to “+” for either flipper tag variable (*s*_2_, *s*_3_) were fixed to zero. The tag loss portion of the state transition generally follows the tag loss model proposed by Laake et al. (2014). Note that the structural zeros only affect the r^(2,3)^ → s^(2,3)^ transitions and there is no interaction term for s^(1)^ and s^(2,3)^; therefore, we can write 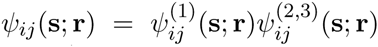. Table A.3 in Supplement A (Johnson et al., 2016a) illustrates the transition probabilities for each current state, r, to each possible state, s.

The full model was fitted with maximum likelihood methods using the forward HMM algorithm in Section 2.3. The ability to fit multivariate state models was added to the R package `marked` (Laake, Johnson and Conn, 2013). R code used for this example is available in Supplement B (Johnson et al., 2016b). Confidence intervals (CI) for derived quantities were approximated using a multivariate normal parametric bootstrap with mean equal to the maximum likelihood estimate and covariance matrix equal to the negative Hessian of the log-likelihood function (see Supplement B; Johnson et al. 2016b). The delta-method could be used. However, with complex parameter functions, forming the gradient can be challenging, thus we opted for the ease of the bootstrap.

### 3.2 Results

The 1996 cohort of sea lion pups on SMI experienced a very strong El Niño in 1997 which both influenced the movement of young animals to ANI and created a high level of mortality before the sea lions reached age 2. With a single cohort and a high level of early mortality, the sample sizes were small at older age classes and the confidence intervals became very wide and that should be considered in evaluating any patterns across age. This unusual event may have also influenced differences in survival across sex and age. However, as an example, it does illustrate the usefulness and flexibility of building multivariate state models.

We begin the results description with the observation model, 5. This is a somewhat unusual example with regard to observation of each state’s status. Observers use the brand for resighting and the tag status was often not observed. Observers were more likely to record status of left tag because the brand is on the sea lion’s left side (Figure 1). The odds of missing second side when the first is missed increases 5-fold [3.5-8.2] (95% confidence intervals are contained within square brackets). This again is due to use of brand for resighting. If the observer didn’t record the status of the left side they were unlikely to get the status of the right side. The fitted detection model, *p*, is illustrated in Figure A.1 of Supplement A (Johnson et al., 2016a). It is relatively standard with respect to multistate (and CJS) modeling, so, we do not elaborate further here.

**Fig 1.**
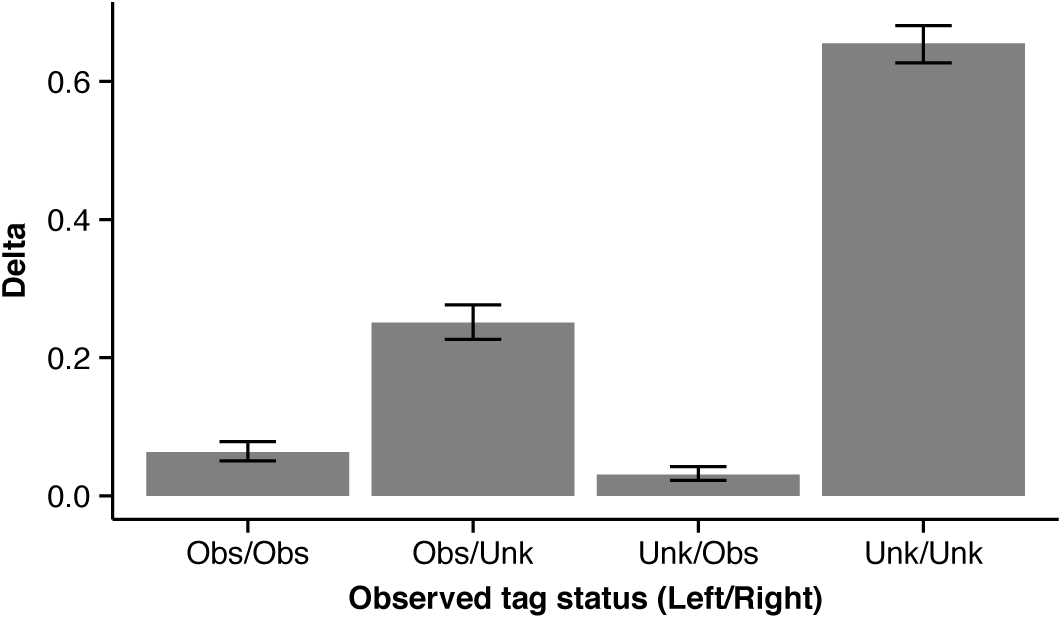
*Estimated δ parameters for the California sea lion model. If the tag status was observed it appears as “Obs” in the plot. Thus, for example, the column “Obs/Unk” indicates the probability that the left tag was observed and the right tag was not observed and its status is unknown. The error bars represent 95% CIs.*

Male pups had a slightly better overall pup survival (Figure 2). Although confidence intervals overlapped considerably. For both sexes, pup survival increased with increasing mass which has also been observed in northern fur seals (Baker and Fowler, 1992; Baker, Fowler and Antonelis, 1994), Steller sea lions (Hastings, Gelatt and King, 2009), and Hawaiian monk seals (Baker, Fowler and Antonelis, 1994; Craig and Ragen, 1999).

**Fig 2.**
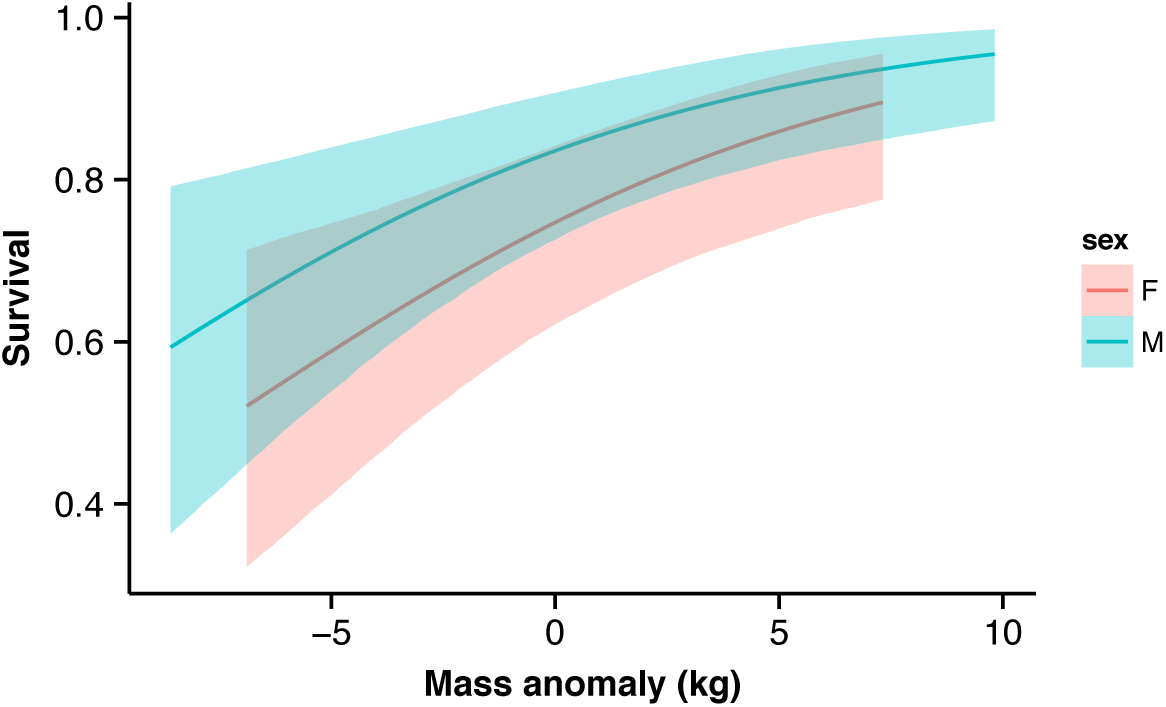
*Mass- and sex-specific survival probability for California sea lion pups on San Miguel Island, California. Mass anomaly is the deviation from the sex-specific mean mass. Colored envelopes represent 95% CIs.*

For non-pups (age > 0) survival increased to adult age (about 5 years old) and then declined in older ages (Figure 3). The decline was much more dramatic for males whose cost of reproduction is high due to their polygynous breeding system in which breeding males have to defend a territory (Peterson and Bartholomew, 1967; Johnson, 1968). For both sexes, higher survival at ANI was likely due to two factors. First, in El Niño years prey resources are distributed farther north, closer to ANI, thus animals not constrained by reproductive commitments at SMI had better access to food. Second, animals at ANI during the breeding season were likely non-reproductive (at least for a given year) and, thus, did not have to invest in costly activities such as holding territories (males) or lactation (females), which allowed them to preserve or invest in better body condition.

**Fig 3.**
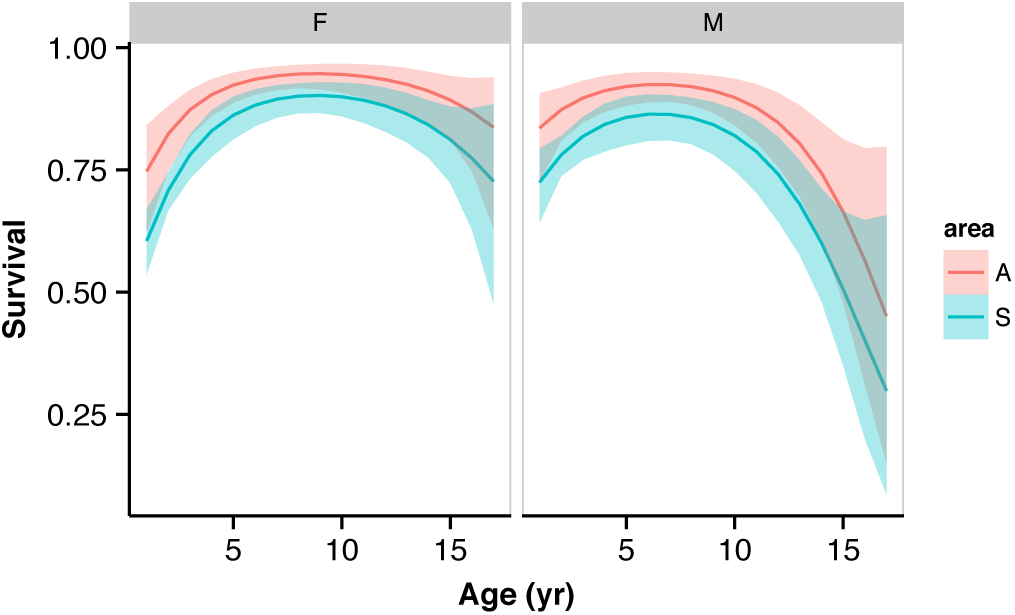
*Sex- and age-specific annual survival probabilities for California sea lion nonpups (age > 0). Separate curves are plotted for each area A = Año Nuevo I. and S = San Miguel I. Although plotted as a solid curve, animals transitioned between the curves depending on the area occupied. Colored envelopes represent 95% CIs.*

With respect to movement between SMI and ANI, we could look at the fitted age and sex-specific annual area transition probabilities (Figure A.2; Supplement A, Johnson et al. 2016b). However, a more biologically interesting, derived quantity is the sex specific proportion of animals that are located in each area for a given age, 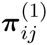. This is because animals at SMI during the breeding season were likely reproductive. This derived quantity can be calculated by

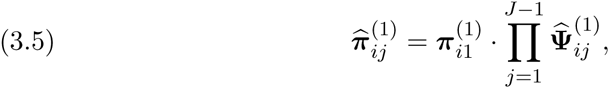

where, for the sea lion data, π*_i_*_1_ = (1, 0)′ because all pups began at SMI on occasion 1, and 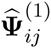 is a 2 × 2 matrix with 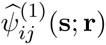 entries. Figure 4A illustrates the occupancy rate of SMI by both sexes. In general, females began returning to SMI at around age 3 and occupancy stabilized to ≈ 0.7 by age 5. Thus, about 70% of females from the 1996 cohort were reproductively active each year based on location alone. This is similar to the 0.77 average natality estimated by Melin et al. (2011) for several cohorts. A large proportion of males stayed at ANI until the age of 10 or 11 when they began to return and to hold territories. Because a large proportion of males were located at ANI during the breeding season, they experienced better annual survival (for the reasons discussed previously) and hence, had higher survivorship than females (Figure 4B). By age 12-13, however, the lower survival for breeding males on SMI resulted in equal survivorship.

**Fig 4.**
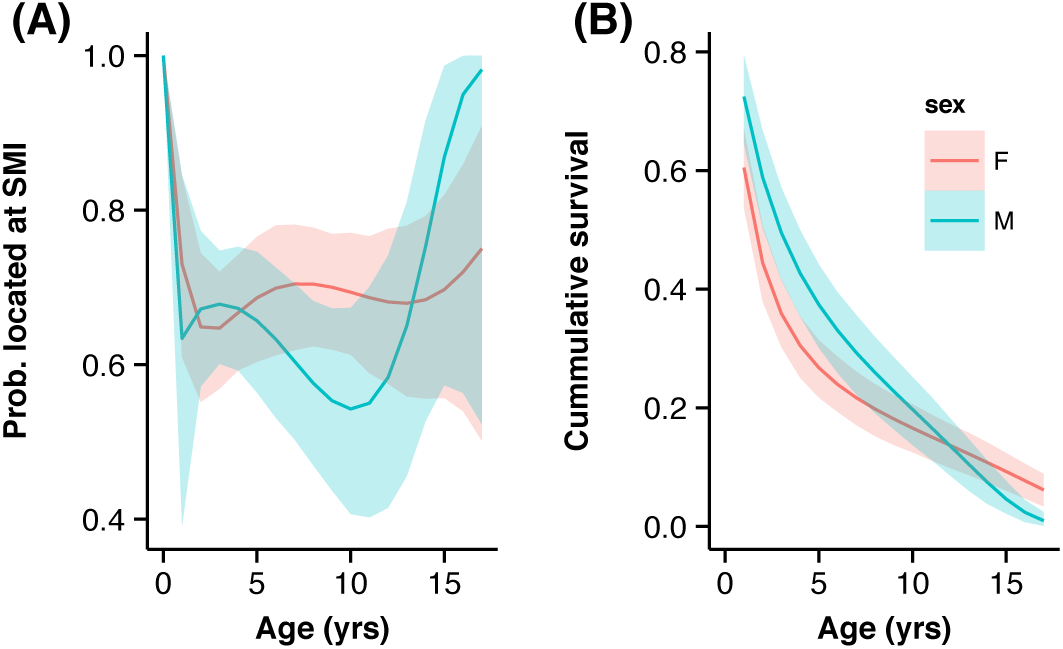
*(A) Age- and sex-specific probability that an individual was located at San Miguel I. during the breeding season. Solid lines and envelopes represent probability that an animal was located, at SMI and 95% CIs. (B) The Age- and sex-specific cumulative marginal survival, i.e., the expected proportion of animals that were alive at a given age.*

Finally, although it did not affect the estimation of survival in this case due to the permanent brand, the flipper tag results were quite revealing for pinniped studies where a third mark is not possible. First, tag loss rate increased with age. Marginally, the odds of loosing a specific tag (left or right) increased 1.56-fold [1.38-1.75] every year. There was also substantial dependence between tags. The odds of losing a tag increased 406-fold [76-2,204] when the other had been lost. Therefore, tags were almost always lost simultaneously. The probability of losing both tags at or before the animal was 5 years old was 0.49 [0.38-0.61]; by age 9, the probability increased to 0.91 [0.86-0.95] (Figure 5; CDF). The most probable age for an animal to become (−, −) tag status was 5-6, when approximately 40% of animals entered double loss status (Figure 5; PDF). This was probably due to the fact that females and males are in periods of strong growth and development at these ages as they prepare for reproductive activities.

**Fig 5.**
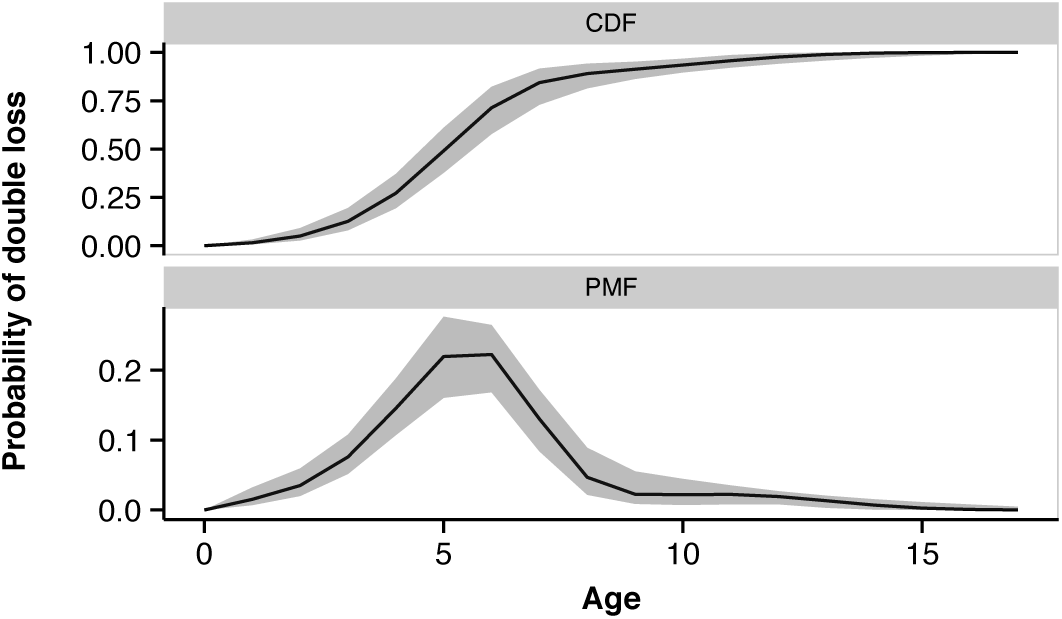
*Age-specific probabilities of double tag loss. The upper plot (CDF) is the cumulative probability distribution of entering the double tag loss state at or before each age. The lower plot (PMF) is the difference of the upper plot and provides the probability mass function that an animal enters the double tag loss state at each age. Gray envelopes represent 95% CIs.*

## 4. DISCUSSION

In the on-going development of state-based CJS modeling we present a generalization of the multievent CJS models to the case where states are multivariate vectors of categorical variables. This allows the user to analyze complex state transitions and handle partial knowledge of states upon observation of the individual. Using a log-linear model parameterization for multiway contingency table data allows parsimonious modeling of multivariate state transitions by enforcing desired independence (or dependence) between state variables by exclusion (inclusion) of specified interaction terms. Because of the necessary and sufficient conditions for determining independence from log-linear models, inference on dependence of state variables can also be tested by examining for significant deviation from zero for the appropriate interaction terms. We use the term “testing” in a loose sense, in that one might ascertain those effects through a model selection procedure (Burnham and Anderson, 2002; Hooten and Hobbs, 2015). Because, we can efficiently calculate the likelihood for the multivariate state model, all of the usual information criteria and Bayesian methods for model selection are applicable.

The parsimonious construction and hypothesis testing properties were both demonstrated in the example analysis of the sea lion data. We assumed that tag loss was primarily due to growth of the foreflippers from the time the tags were placed on the animals as pups. Therefore, it is reasonable to conclude that loss of tags will be independent of the location of the animal (i.e., tag loss variables *s*_2_ and *s*_3_ are independent of the area *s*_1_). Therefore, we specifically omitted any interaction terms that would imply dependence of tag loss on area. Second, we examined the parameter measuring interaction between the loss of each tag, *β*^(2,3)^(s; r), for significant deviation from zero and found that there is a high degree of dependence in the loss of flipper tags in California sea lions. This conclusion has strong implications for future studies that do not use a permanent third mark such as a brand, as survival estimation may be negatively biased, depending on the assumed tag loss process (Laake et al., 2014). By using the multivariate state framework we were able to directly extend the double tag loss model of Laake et al. (2014) to account for movement between different areas as well. Another practical extension multivariate models provide over the single multievent framework is that the probability of state uncertainty can depend on the true state. For example, although we did not examine it here, it is possible that the probability of an observer missing the status of a flipper tag depends on whether it is present or not. Or, if there are different observer teams in each area, one team may ignore the tag status, while the other dutifully records it. Thus, probability of correct observation of one state variable may depend on another state variable.

Although the proposed multivariate framework is quite general, there are further extensions that are straightforward to develop given the HMM formulation. First, our model was based on the assumption that the state variable is either observed correctly or it is not observed at all and hence, unknown. There is no reason that this cannot be relaxed. All that needs to be done is to augment the observation space for each variable for which errors can be made. For example, 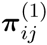, where *e*_1_, … refers to states that are not biological in origin, but observational errors. In the tag loss example, an observer might see a tag is physically present, but fail to read it. In this case it is uncertain whether or not the tag should be considered lost (i.e., unreadable = lost), but there is more information than a simple “unknown” observation. In addition, it is also possible that an observer might declare a tag to be lost when in fact it is present or *vice versa.* This is accomplished by removing the constraint that *δ_ij_*(c; s) = 0 when any *c_k_* ≠ *s_k_* or *u*. Finally, continuously valued states could be incorporated using the discrete approximation method of Langrock and King (2013). This creates a large state-space for the continuous variable, thus, careful consideration of the interaction parameterization may be necessary. Each of these three extensions are directly handled by the HMM inference framework. Some of these extensions, however, might produce a model of such complexity that some parameters become unidentifiable or the model fitting is computationally unfeasible. Placement of the multivariate model in a robust design framework similar to Kendall et al. (2012) might improve estimation of parameters, however, careful parameterization may be the only way to handle large state-spaces. Robust design analysis can be accomplished in the multivariate setting (and in `marked` package in general) by constraining parameters to be constant (most likely 0 or 1) within secondary sampling periods, but free between primary periods.

In addition to model extensions there is also need to develop methods to assess goodness-of-fit (GOF) for complex CJS models such as those presented here. We do not provide specific recommendations here, as this is a large topic in its own right. There are, however, some avenues worth pursuing. Zucchini and MacDonald (2009) and Titman and Sharples (2008) provide general frameworks for assessing model fit in HMMs. Those general approaches may be adapted for the specifics of the state-based CJS models. In addition, the minimally sufficient statistics GOF testing approach of Pradel, Gimenez and Lebreton (2005) might be investigated using the work of King and McCrea (2014) to derive the necessary sufficient statistics.

We, again, would like to acknowledge the seminal work by the Cormack (1964), Jolly (1965), and Seber (1965) papers which initiated 50 years of development and analysis of capture-recapture data. Through the years, the same basic framework has been generalized to add additional inference capability as well as overcome assumptions of the original CJS models, but the same basic framework exists in each of these models, including the extension proposed herein.

## ACKNOWLEDGMENTS

The findings and conclusions in the paper are those of the authors and do not necessarily represent the views of the National Marine Fisheries Service, NOAA. Reference to trade names does not imply endorsement by the National Marine Fisheries Service, NOAA.

## SUPPLEMENTARY MATERIAL

### Supplement A: Notation summary and additional sea lion analysis details

(doi: 10.1214/00-AOASXXXXSUPP; supplement_A.pdf). Contains additional tables that summarize notation used throughout the paper and provide additional details and results for the analysis of sea lion data in Section 3.

### Supplement B: R code used to analyze sea lion data

(doi: 10.1214/00-AOASXXXXSUPP; supplement_B.pdf). Contains the R code used to run the sea lion example analysis in Section 3. Versions ≥ 1.1.10 of `marked` contain the multivariate state fitting capability and the sea lion data. Using the R console, the following command will install the package:

~~~
install.packages("marked")
~~~

a precompiled version can be installed from CRAN.

## Supplement A: Notation summary and additional example results and details

### Supplement to “Multivariate state hidden Markov models for mark-recapture data”

#### Acknowledgments

**Table A.1.**
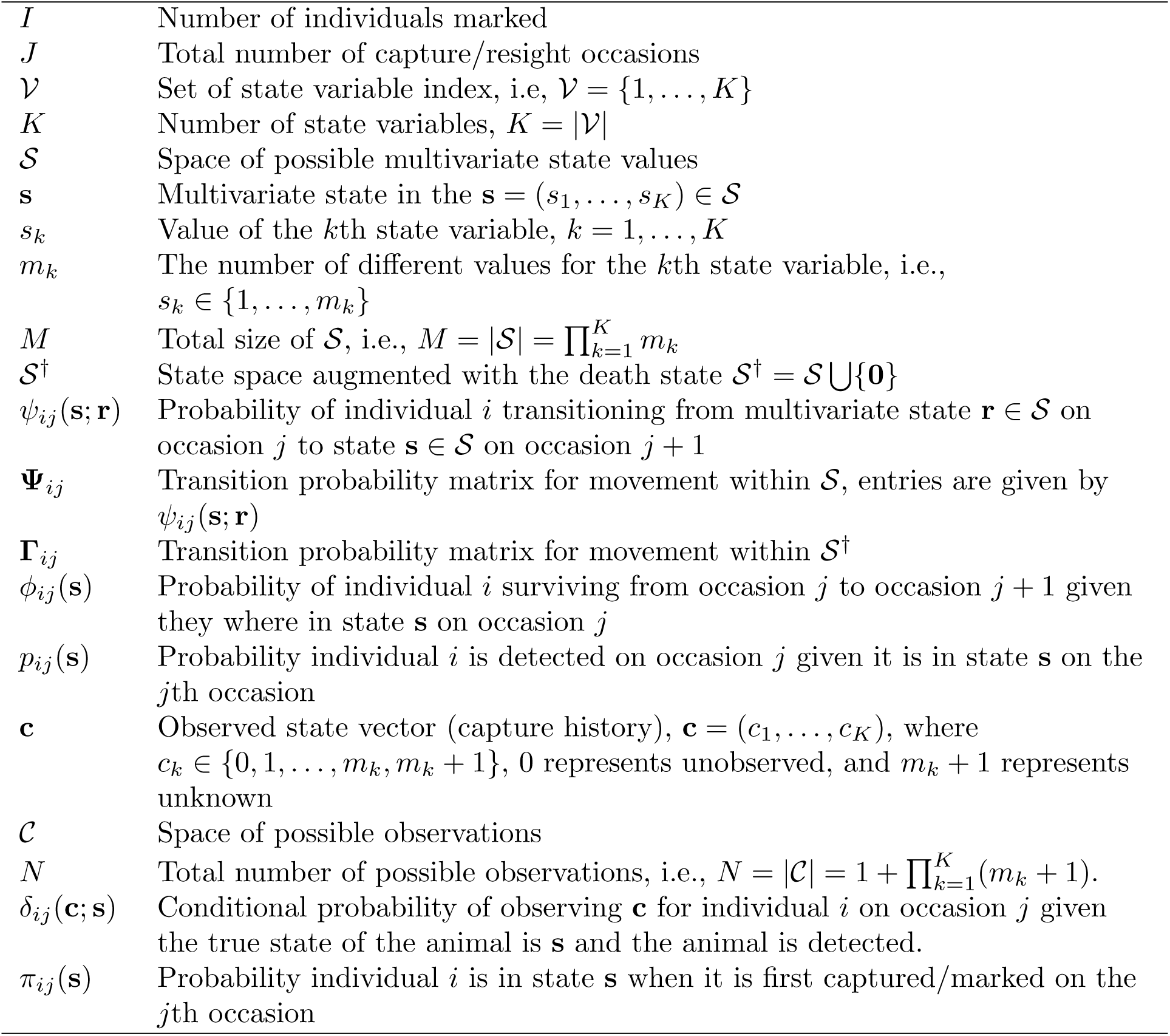
*Notation glossery. Here we present a list of notation used throughout the paper. The individual index I runs from 1 to n, the total number of marked individuals and j runs from 1 to T, the total number of capture/resighting occasions.*

**Table A.2.** *Conditional probabilities, δ*(c; s), *for the observations,* c *given the true state,* s*, of detected, sea lion in the example analysis of Section 3. Because the location, s*_1_, *is known with certainty when the animal is detected, we can write δ*(c; s) = *δ*(c^(2,3)^; s^(2,3)^). *Here we use the true state and the reference cell. Structural zeros occur for observations where the state is mis-observed rather that just unobserved. The δ probabilities are presented in their unnormalized form. The necessary normalizing constants are the sums over each column. The value in the last row holds for all columns.*

| | | $\mathbf{s}^{(2,3)}$ | | | |
| --- | --- | --- | --- | --- | --- |
| | | $(+, +)$ | $(+, -)$ | $(-, +)$ | $(-, -)$ |
| $\mathbf{c}^{(2,3)}$ | $(+, +)$ | 1 | 0 | 0 | 0 |
| | $(+, -)$ | 0 | 1 | 0 | 0 |
| | $(-, +)$ | 0 | 0 | 1 | 0 |
| | $(-, -)$ | 0 | 0 | 0 | 1 |
| | $(+, u)$ | $\exp\{\alpha^{(3)}\}$ | $\exp\{\alpha_3\}$ | 0 | 0 |
| | $(-, u)$ | 0 | 0 | $\exp\{\alpha^{(3)}\}$ | $\exp\{\alpha^{(3)}\}$ |
| | $(u, +)$ | $\exp\{\alpha^{(2)}\}$ | 0 | $\exp\{\alpha^{(2)}\}$ | 0 |
| | $(u, -)$ | 0 | $\exp\{\alpha^{(2)}\}$ | 0 | $\exp\{\alpha^{(2)}\}$ |
| | $(u, u)$ | $\exp\{\alpha^{(2)} + \alpha^{(3)} + \alpha^{(2,3)}\}$ | | | |

**Table A.3a.** *Transition probabilities,* 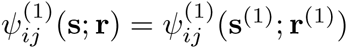, *for the movement between areas, s*_1_ = *s (San Miguel I.) and s*_1_ = *a (Año Nuevo I.). Readers should recall that* 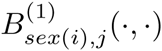 *is a sex-specific, 3 df b-spline smooth over age = j* − 1*. Probabilities are formed by normalizing over rows of the table entries.*

| | | $\mathbf{s}^{(1)}$ | |
| --- | --- | --- | --- |
| | | $s$ | $a$ |
| $\mathbf{r}^{(1)}$ | $s$ | 1 | $\exp\{\beta_{sex(i)}^{(1)}(a; s) + B_{sex(i),j}^{(1)}(a; s)\}$ |
| | $a$ | $\exp\{\beta_{sex(i)}^{(1)}(s; a) + B_{sex(i),j}^{(1)}(s; a)\}$ | 1 |

**Table A.3b.**
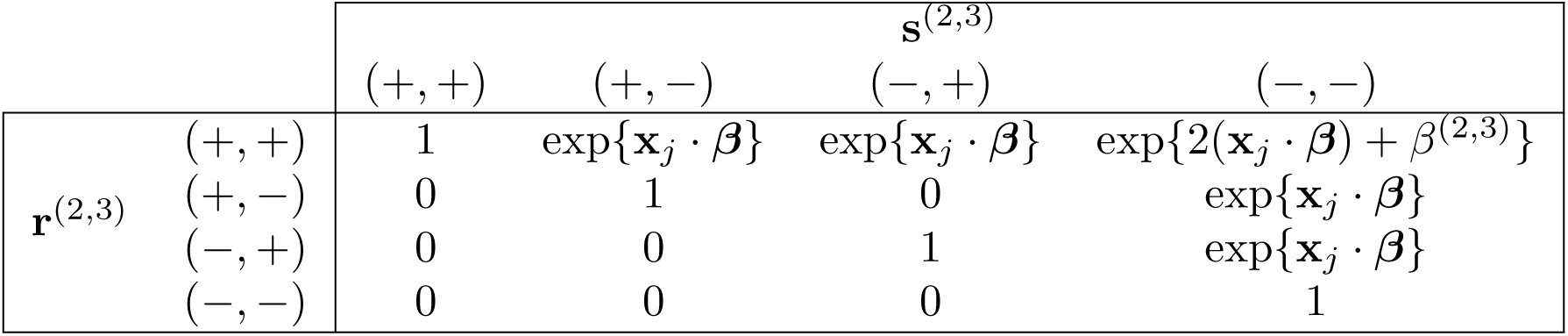
*Transition probabilities,* 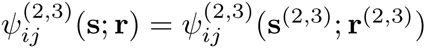, *, for the tag loss model presented in Section 3. The state of the K* − 1*th flipper tag, s_k_*, *k* = 2, 3 *is represented by s_k_* ∈ {+, −}, *where a “+” indicates the tag is present and “−” indicates the tag has been lost. Recall from the main portion of the paper that because the tags are identical, we assumed that the marginal loss rate would be the same for each side. Therefore, we constrained the non-zero parameter values β*^(2)^( −;+) = *β*^(3)^(−;+) = *β. In addition, we modeled a linear effect of age, so the covariate vector* x*_j_* = (1, *j* − 1). *Values of zero, are due to the fact that tag loss is an absorbing state from which the animal cannot return. The ψ probabilities are presented in their unnormalized form, necessary normalizing constants are the sums over each row.*

| | | $\mathbf{s}^{(2,3)}$ | | | |
| --- | --- | --- | --- | --- | --- |
| | | $(+, +)$ | $(+, -)$ | $(-, +)$ | $(-, -)$ |
| $\mathbf{r}^{(2,3)}$ | $(+, +)$ | 1 | $\exp\{\mathbf{x}_j \cdot \beta\}$ | $\exp\{\mathbf{x}_j \cdot \beta\}$ | $\exp\{2(\mathbf{x}_j \cdot \beta) + \beta^{(2,3)}\}$ |
| | $(+, -)$ | 0 | 1 | 0 | $\exp\{\mathbf{x}_j \cdot \beta\}$ |
| | $(-, +)$ | 0 | 0 | 1 | $\exp\{\mathbf{x}_j \cdot \beta\}$ |
| | $(-, -)$ | 0 | 0 | 0 | 1 |

**Fig. A.1.**
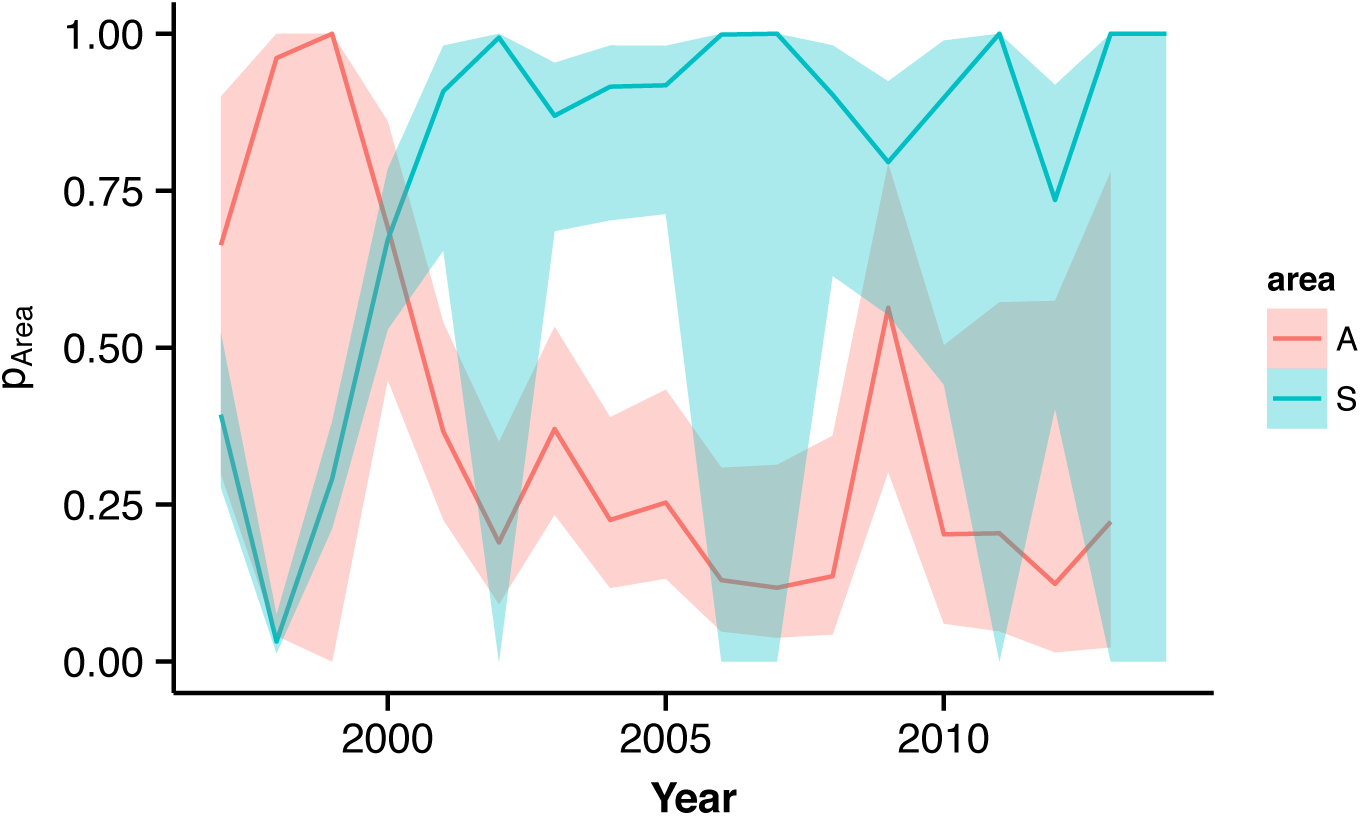
*Area- and occasion-specific detection probabilities for California sea lions.*

**Fig. A.2.**
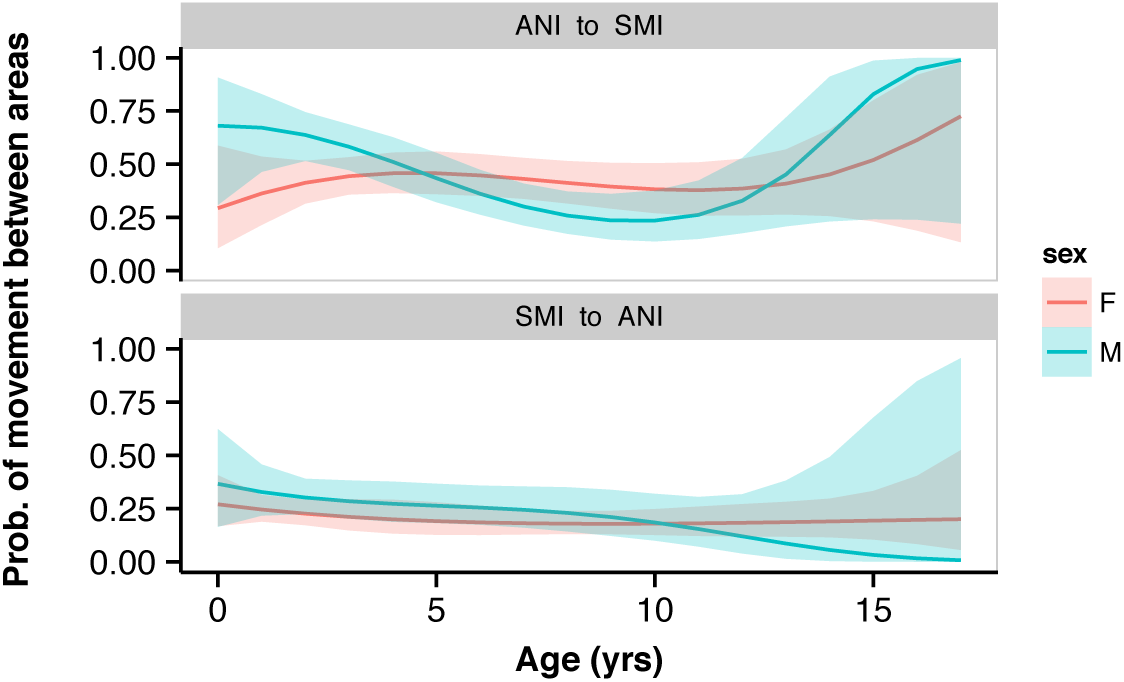
*Sex-specific probability of movement between San Miguel Island and Año Nuevo Island, California*

## Supplement B: R code used to analyze sea lion data

### Supplement to “Multivariate state hidden Markov models for mark-recapture data”

#### Acknowledgments

Data analysis with the multivariate state models can be accomplished with the marked package but a certain level of proficiency with R programming is required. In addition, a thorough knowledge of modeling mark-recapture data and use of multinomial logit link function is needed. Here we provide the code used to fit the model to the sea lion data as an example. It can be used as a template for other analysis. Laake et al (2013) provide an overview of the marked package which will suffice as an introduction but here we describe the extensions that enable multivariate state models to be fitted with the package.

First we attach the marked package and additional packages used for analysis, prediction and graphics.

~~~
### Load packages ###
# *Must have 'marked' v >= 1.1.10*
library (marked)
# *The splines package is only necessary for fitting b-spline curves used in the paper*
# *It is not required for the multivate models in the marked package*
library (splines)
library (mvtnorm)
library (dplyr)
library (ggplot2)
library (cowplot)
~~~

The example data are provided in the marked package and can be retrieved with `data(sealions)`. The data contain a capture history string, a numeric weight value which is the anomaly from the sex-specific mean in kg and a sex value (`F` or `M`) for 485 sea lions. The sea lions were initially marked and tagged as pups in 1996 on San Miguel Island, CA. The capture history string has a 3-character representation for each occasion and the occasion values are separated by a comma. The first character is either `A` (Año Nuevo Island) or `S` (San Miguel Island) for the animals location when it was seen. The second and third characters represent the status of the left and right tags respectively. A “+” means the tag was present, a means the tag was absent and a “−” means the tag status was not observed (unknown).

~~~
data(sealions)
~~~

The first step in using the marked package is to process the data. Processing the data includes specifying mark-recapture model to be used (mvmscjs) and additional data attributes (e.g., strata.labels). For this model, the labels and values for the capture history string must be provided from left to right in the string. The following specifies that the first variable is called “area” and the possible characters are `“A”` and `“S”` as described above. The second variables are named `ltag` and `rtag` and have values `“+”`,`“−”`, or `“u”`. The tag status variables can be unknown but the area is always known because a `“u”` was not included in the vector for area.

~~~
# *Process data for multivariate models in marked*
dp=process.data(sealions,model="mvmscjs",                   strata.labels=list(area=c("A","S"),ltag=c("+","−","u"),rtag=c("+","−","u")))
## 485 capture histories collapsed into 334
~~~

In processing the data, if there are duplicate data records they are collapsed into unique records (referenced by `id`) and a field named `freq` contains the number of sea lions with the same data. In this case the 485 records are collapsed to *I* = 334 ids with frequencies ranging from 1 to 16. The most commonly collapsed records are those that are released and never seen again. Collapsing into the unique records reduces execution time and the size of the design data list.

The processed data list (`dp`) contains the data and the model and data attributes. The processed data list is passed to the function `make.design.data` which creates a list of design dataframes with a dataframe for each parameter. The parameters for the “`mvmscjs`” models are `Phi, p, Psi` and `delta` for this model. It does not include `pi` because this implementation assumes that the variables are all known at the time of release.

~~~
### Make design data
ddl=make.design.data(dp)
~~~

The dataframe for `Phi` contains a record for each id-interval-stratum where stratum is all *M* combinations of the values of the variables used to define the states. For this example there are 8 strata (A, A+−, A−+, A−, S, S+−, S−+, S−) and 19 occasions and 18 intervals. Thus, there are *I* (*J* − 1)*M* = **18** × 8 = *I* × 144 records in the design data. Likewise, for `p` there is a record for id-occasion-stratum (I(*J −* 1)*M* = *I* × 144) where occasions are resight occasions from 2 to 19. A logit link function is used for `Phi` and `p`.

~~~
names(ddl$Phi)
~~~

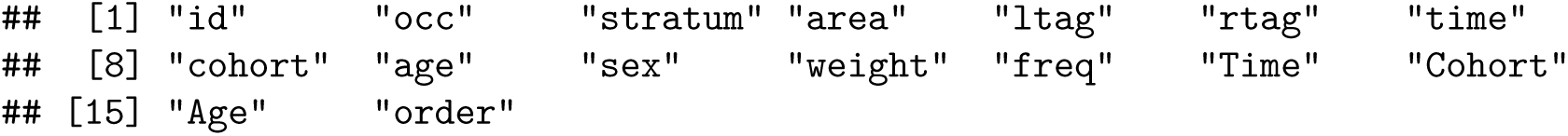

~~~
nrow(ddl$Phi[ddl$Phi$id==1,])
~~~

\## [1] 144

~~~
names(ddl$p)
~~~

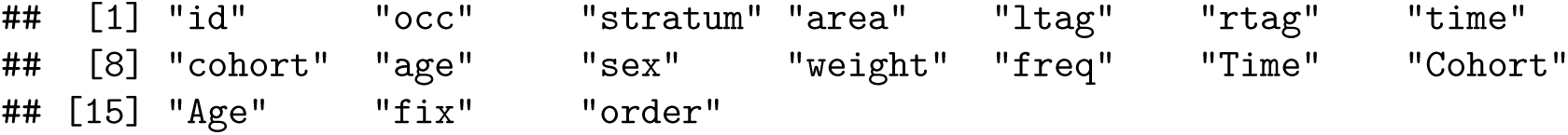

~~~
nrow(ddl$p[ddl$p$id==1,])
~~~

\## [1] 144

For the `delta` design data, there are *I (J − 1)Mn* records where *n* is the possible number of observations for each state. In this example, *n* = 4 because you can either observe the state variable or unknown value for two of the three variables, `ltag` and `rtag`. For example, for state `A++` you can observe `A++, A+u, Au+`, or `Auu`. Each set of *n* records is a multinomial with the sum of the four probabilities equal to 1. The multinomial logit is implemented using a log link function and then normalizing by the sum over the set of *n* values. To be identifiable, one of the *n* values must be fixed to 1 (reference cell). We will explain fixed values later. As a convention we suggest specifying the fully known observation (e.g., `A++`) as the reference cell. Had we specified no `u` values for the variables then there would have only been *n* = 1 record which would be fixed to 1 (i.e., no unknown values). For each id and occasion, there are *Mn* = 8 × 4 = 32 records.

~~~
head(ddl$delta)
~~~

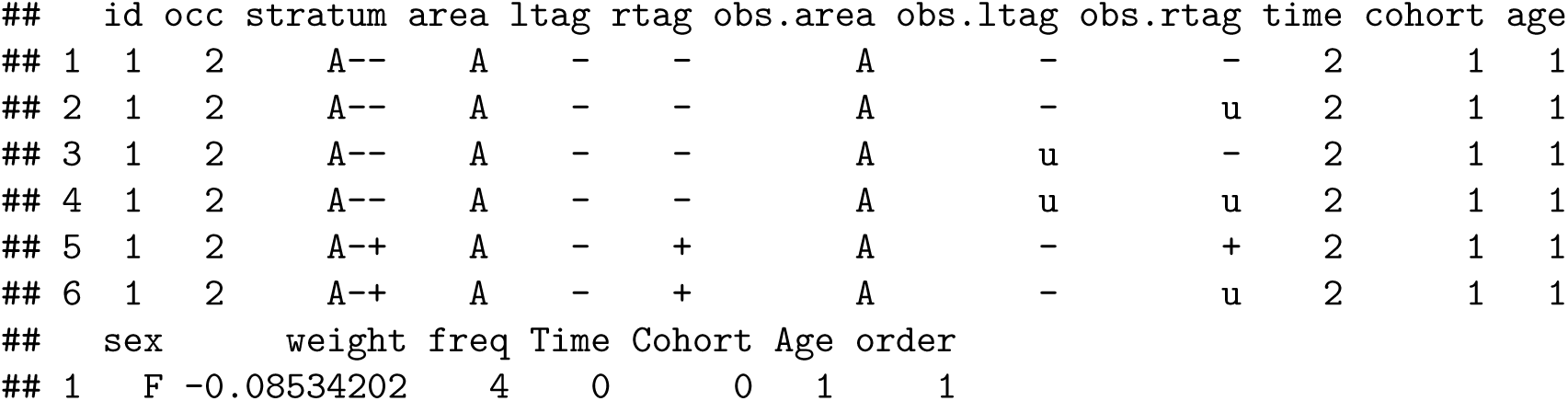

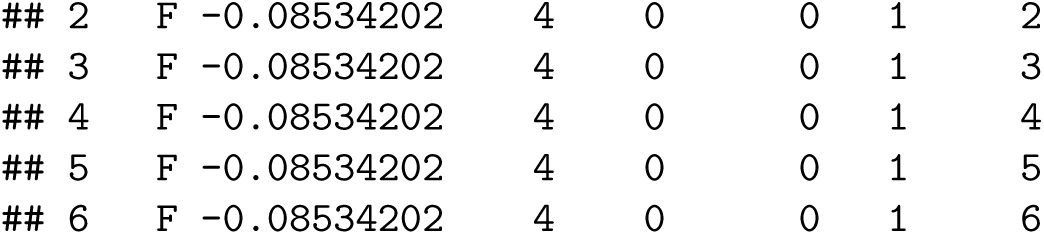

~~~
nrow(ddl$delta[ddl$delta$id==1&ddl$delta$occ==2,])
~~~

\## [1] 32

For the Psi design data, there are *I (J −* 1)*M*_2_ = *I* × 18 × 64 = *I* × 1152 records. For each id-occasion-state, there are a set of *M* records that represent a multinomial for the probability of transitioning from that state to any of the *M* states. The reference cell is set up by default to be Ψ*_rr_* of staying in the same state (r → r). This is done when the design data are created by adding the field “`fix`” and assigning its value to `NA` except for the reference cell which get the value 1. Values with `NA` are estimated and any other value fixes the real parameter (inverse link) at that value. The value 1 is the real parameter value for a log link with a beta value of 0 (1=exp(0)). The multinomial parameters, `delta` and `Psi` are set up with a log link but then normalized by the sum over the multinomial set. For Psi that is a set of *M* records with 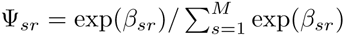 and exp(*β_rr_*) = 1. A different reference cell could be used by changing the values of the field `fix`. The reference cell does not have to be the same for each state or it could vary by id or occasion as well but specifying the formula could become tricky in the latter cases.

~~~
head(ddl$Psi)
~~~

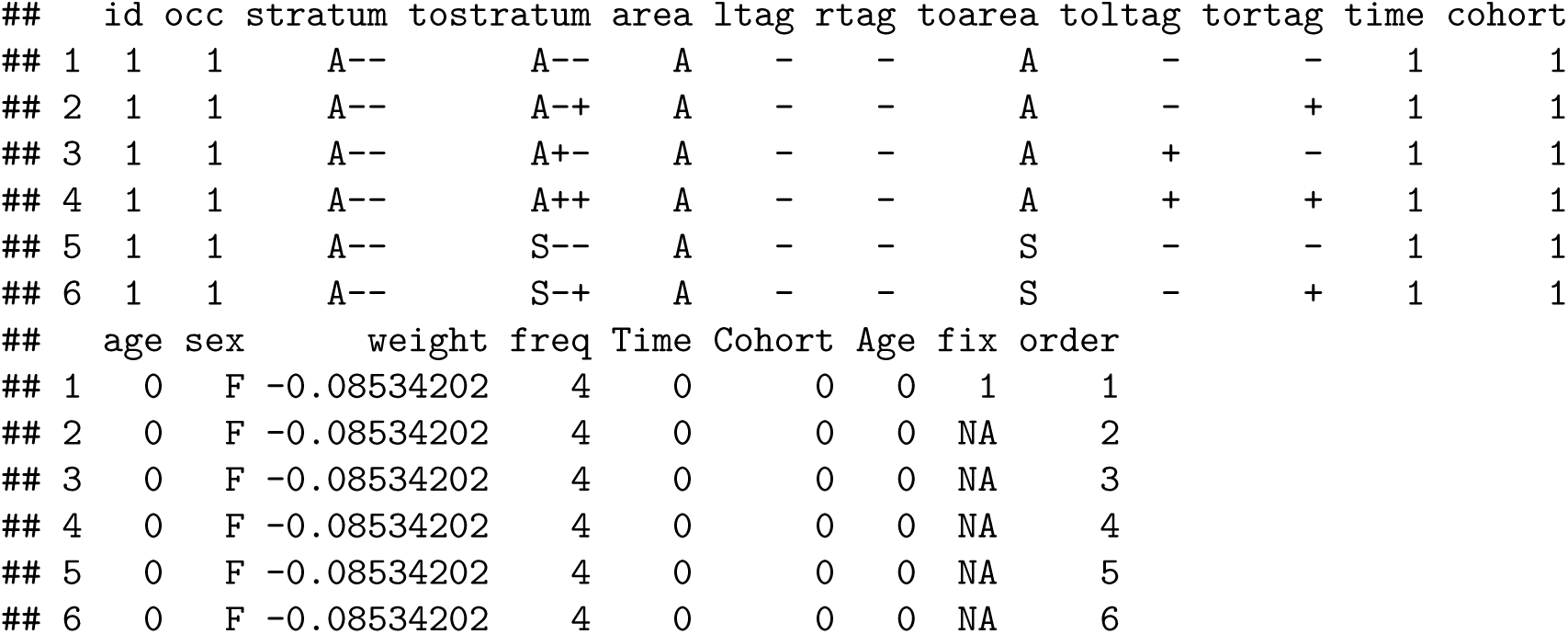

~~~
nrow(ddl$Psi[ddl$Psi$id==1&ddl$Psi$occ==1,])
~~~

\## [1] 64

Default design data are created by `make.design.data` but you are free to add your own fields which may create other fields derived from the default fields or possibly add entirely new data fields by merging in occasion data like environmental variables. If you do merge in other data with the merge function, make sure to re-order the design dataframe using the variable order. Below, a variable pup is defined for `Phi` and `sex` is changed from a character to a factor variable. Had the latter been done in the data, this would not have been needed.

~~~
# *Create pup variable for Phi*
ddl$Phi$pup=ifelse(ddl$Phi$Age==0, 1, 0)
ddl$Phi$sex=factor(ddl$Phi$sex)
~~~

Data from Año Nuevo Island in 2014 were not available at the time, so `p` is set to `0` for `area=="A"` on the last occasion:

~~~
# *Detection model*
# *Set final year p=0 (no resight data) for ANI*
ddl$p$fix = ifelse(ddl$p$Time==17 & ddl$p$area=="A", 0, NA)
~~~

For `delta`, a 0/1 variable is created for the left and right tag when it is unknown. These variables are used in the formula such that the implied reference cell is when the status of both tags is known.

~~~
# *Delta model*
# *create indicator variables for 'unknown' tag observations*
ddl$delta$obs.ltag.u = ifelse(ddl$delta$obs.ltag=="u", 1, 0)
ddl$delta$obs.rtag.u = ifelse(ddl$delta$obs.rtag=="u", 1, 0)
~~~

The changes to the `Psi` data are slightly more involved. We cannot have transitions from “`−`” (missing) to “+” present for a tag, so that transition is set to `0` for both the left and right tag. Dummy 0/1 variables are created for transitions from `A` to `S` and from `S` to `A` to be used in the formulas for `Psi`. In addition, dummy variable `lpm` and `rpm` are set to 1 for transitions from “+” to “`−`” (losing a tag) and 0 otherwise. Finally, `sex` is converted to a factor variable.

~~~
# *Psi model*
# *Set Psi to 0 for cases which are not possible - missing tag to having tag*
ddl$Psi$fix[as.character(ddl$Psi$ltag)=="-"&as.character(ddl$Psi$toltag)=="+"]=0
ddl$Psi$fix[as.character(ddl$Psi$rtag)=="-"&as.character(ddl$Psi$tortag)=="+"]=0
# *Create indicator variables for transitioning between states*
ddl$Psi$AtoS=ifelse(ddl$Psi$area=="A"&ddl$Psi$toarea=="S",1,0) *# ANI to SMI movement*
ddl$Psi$StoA=ifelse(ddl$Psi$area=="S"&ddl$Psi$toarea=="A",1,0) *# SMI to ANI movement*
ddl$Psi$lpm=ifelse(ddl$Psi$ltag=="+"&ddl$Psi$toltag=="-",1,0) *# Losing left tag*
ddl$Psi$rpm=ifelse(ddl$Psi$rtag=="+"&ddl$Psi$tortag=="-",1,0) *# Losing right tag*
ddl$Psi$sex=factor(ddl$Psi$sex)
~~~

Formulas are specified in a list and named by the variable with a dot and an extension (eg 1 in each case below). This form is suggested to use the crm.wrapper function for fitting multiple sets of models.

~~~
Psi.1=list(formula=—1+ AtoS:sex + AtoS:sex:bs(Age) + StoA:sex + StoA:sex:bs(Age) + I(lpm+rpm) +I(lpm+rpm):Age + lpm:rpm)
p.1=list(formula=~time*area)
delta.1=list(formula= ~ -1 + obs.ltag.u + obs.rtag.u + obs.ltag.u:obs.rtag.u)
Phi.1=list(formula=~sex*bs(Age)+pup:weight+area)
~~~

The formula for `p` is the simplest. It specifies time varying resighting probabities that differ across the two areas (`A` and `S`). The formula for `Phi` specifies a cubic spline across age that varies by `sex`, survival varying by weight for `pup==1`, and an additive `area` difference in survival. The formula for delta specifies a dependence model for the probability of not observing the status of a tag in which if one tag is not known may influence whether the other tag is known. By removing the intercept (−1), the intercept value is 0 which fixes the reference cell to be observed status for both tags (`obs.ltag.u=0` and `obs.rtag.u=0`). The formula for `Psi` is the most complicated involving two additive portions with the first for movements between `A` and `S` and `S` to `A` that are a function of `sex` and `Age` represented by a spline that is sex specific. The second portion is for tag loss (transitions from “+” to “`−`”). It specifies tag loss that is constant for left and right (I(lpm+rpm)), but changes linearly with age (`I(lpm+rpm):Age`) and dependence (`lpm:rpm`) whereby losing one tag influences loss of the other tag.

For this single model a call to `crm` fits the model and returns the results into `mod`. The arguments are the processed data list (`dp`), the design data (`ddl`), the `model.parameters` which is a list of the parameter lists and an argument hessian set to `TRUE` to get the variance-covariance matrix. Note that this model fitting with estimation of the hessian may take about an hour to run. We are working on finding more efficient ways of fitting these models. With the generality of fitting models in marked, comes the concomitant cost of slow execution.

~~~
# *Fit model*
mod=crm(dp,ddl,model.parameters=list(Psi=Psi.1,p=p.1,delta=delta.1,Phi=Phi.1,hessian=TRUE)
~~~

Various plots were produced from the model results. Here we provide the code to produce plots for survival. The first plot shows the pup survival estimates as a function of their weight anomaly. The comments below describe the code used.

~~~
*#pup - create a dataframe with a sequence of weights for females and males*
Phi_pup = rbind(
   data.frame
      ( sex="F",
      weight=seq(min(ddl$Psi$weight[ddl$Psi$sex=="F"]),
                  max(ddl$Psi$weight[ddl$Psi$sex=="F"]),0.1)
),
data.frame
   ( sex="M",
   weight=seq(min(ddl$Psi$weight[ddl$Psi$sex=="M"]),
                      max(ddl$Psi$weight[ddl$Psi$sex=="M"]),0.1)
)
)
# *create a design matrix for the pup survival with the data defined above*
Phi_pup_dm = model.matrix(~sex + weight, data=Phi_pup)
Phi_coef = mod$results$beta$Phi
# *extract the variance-covariance matrix for the betas*
vcv = mod$results$beta.vcv
vcv_names = rownames(mod$results$beta.vcv)
# *extract the portion of the v-c matrix for the Phi parameters*
Phi_vcv = vcv[grep("Phi", vcv_names),grep("Phi", vcv_names)]
# *compute 10,000 multivariate normal variables with means Phi_coef*
# *and variance-covariance matrix Phi_vcv*
Phi_rep = rmvnorm(10000, Phi_coef, Phi_vcv)
# *using the random variable values for the intercept, sex and weight variables compute the*
# *survival estimates for the 10000 variates*
pred_rep=apply(Phi_rep[,c(1,2,10)], 1, FUN=function(b,D){plogis(D%*%b)}, D=Phi_pup_dm)
# *compute the 95% confidence intervals from the simulated values*
tmp = t(apply(pred_rep, 1, quantile, prob=c(0.025, 0.975)))
colnames(tmp) = c("lower","upper")
# *create data frame for plotting and call ggplot to create the plot*
Phi_pup = cbind(Phi_pup,tmp)
Phi_pup$estimate = plogis(Phi_pup_dm%*%Phi_coef[c(1,2,10)])
p_pup_surv = ggplot(data=Phi_pup) + geom_line(aes(x=weight, y=estimate, color=sex)) + geom_ribbon(aes(ymin=lower, ymax=upper, x=weight, fill=sex), alpha=0.33) + ylab("Survival") + xlab("Mass anomaly (kg)") + geom_rangeframe()
p_pup_surv
~~~

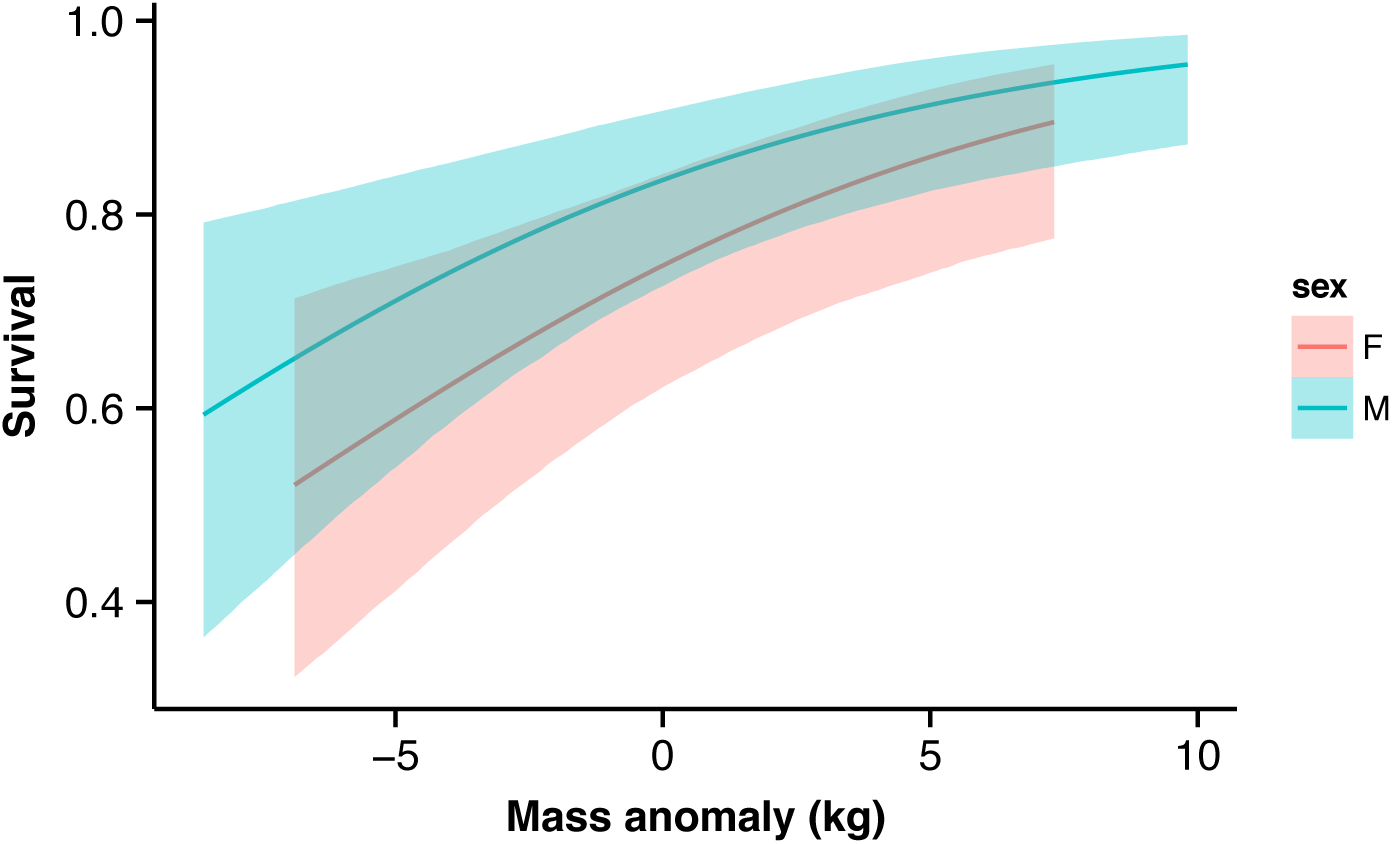

The second plot shows the non-pup survival estimates as a function of age. The procedure used is very similar to the code used above.

~~~
*#adult - Follow a similar*
Phi_ad = data.frame
    ( expand.grid(
        sex=c("F","M"),
        Age=c(1:17),
        area=c("A","S"),
        pup=0,
        weight=0
  )
)
Phi_ad_dm = model.matrix(mod$model.parameters$Phi$formula, Phi_ad)
Phi_ad$estimate = plogis(Phi_ad_dm%*%Phi_coef)
tmp = Phi_rep %>% apply(., 1, FUN=function(b,D){plogis(D%*%b)}, D=Phi_ad_dm) %>% apply(., 1, quantile, prob=c(0.025, 0.975)) %>% t(.)
colnames(tmp)=c("lower","upper")
Phi_ad = cbind(Phi_ad,tmp)
p_ad_surv = ggplot(data=Phi_ad) + geom_line(aes(x=Age, y=estimate, color=area)) + geom_ribbon(aes(ymin=lower, ymax=upper, x=Age, fill=area), alpha=0.33) + facet_wrap(~sex) +
    ylab("Survival") + xlab("Age (yr)")
p_ad_surv
~~~

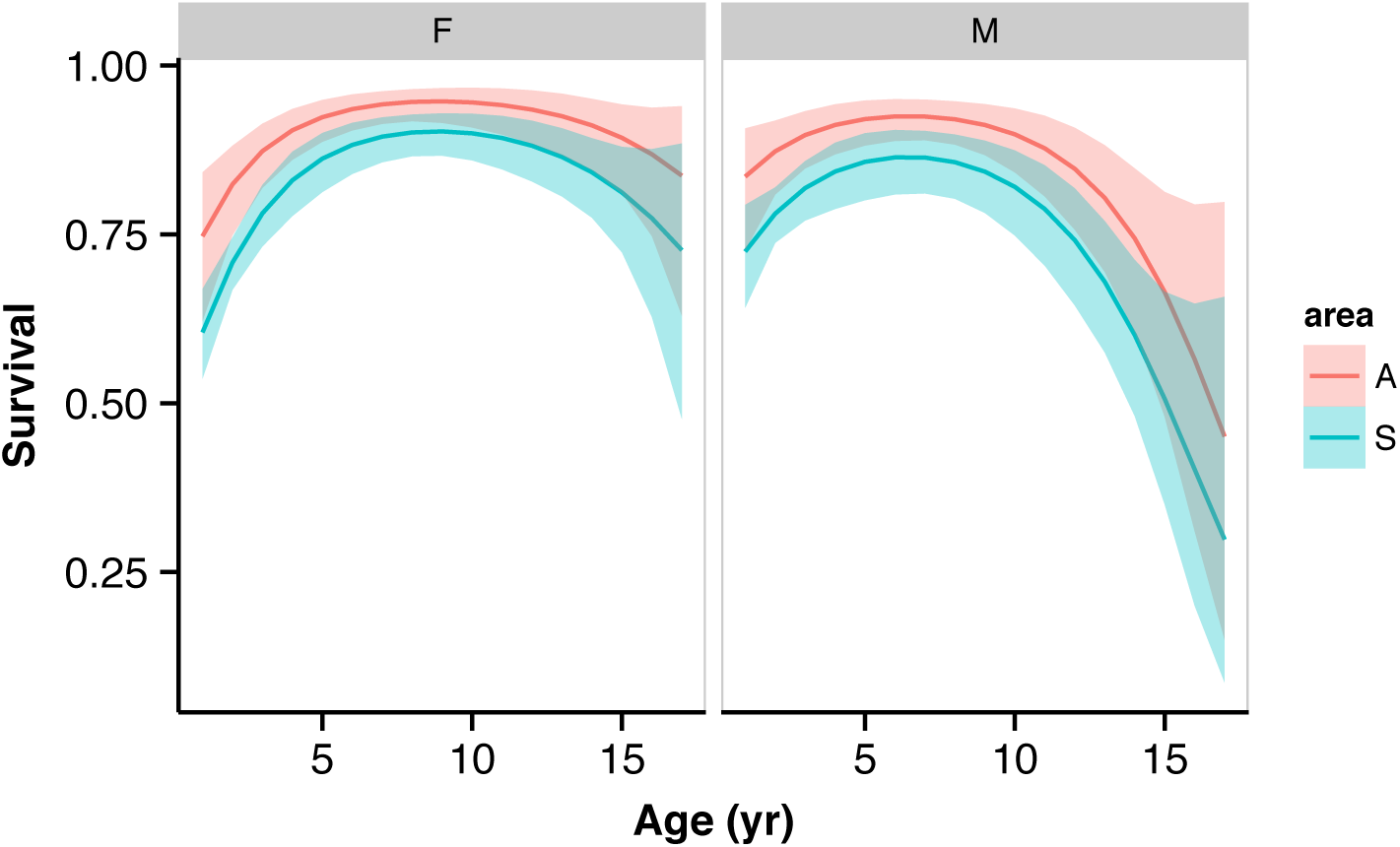

